# Characterization of natural product inhibitors of quorum sensing in *Pseudomonas aeruginosa* reveals competitive inhibition of RhlR by *ortho*-vanillin

**DOI:** 10.1101/2024.02.24.581676

**Authors:** Kathryn E. Woods, Sana Akhter, Blanca Rodriguez, Kade A. Townsend, Nathan Smith, Ben Smith, Alice Wambua, Vaughn Craddock, Rhea G. Abisado-Duque, Emma E. Santa, Daniel E. Manson, Berl R. Oakley, Lynn E. Hancock, Yinglong Miao, Helen E. Blackwell, Josephine R. Chandler

## Abstract

Quorum sensing (QS) is a cell-cell signaling system that enables bacteria to coordinate population density-dependent changes in behavior. This chemical communication pathway is mediated by diffusible *N*-acyl L-homoserine lactone signals and cytoplasmic signal-responsive LuxR-type receptors in Gram-negative bacteria. As many common pathogenic bacteria use QS to regulate virulence, there is significant interest in disrupting QS as a potential therapeutic strategy. Prior studies have implicated the natural products salicylic acid, cinnamaldehyde and other related benzaldehyde derivatives as inhibitors of QS in the opportunistic pathogen *Pseudomonas aeruginosa,* yet we lack an understanding of the mechanisms by which these compounds function. Herein, we evaluate the activity of a set of benzaldehyde derivatives using heterologous reporters of the *P. aeruginosa* LasR and RhlR QS signal receptors. We find that most tested benzaldehyde derivatives can antagonize LasR or RhlR reporter activation at micromolar concentrations, although certain molecules also caused mild growth defects and nonspecific reporter antagonism. Notably, several compounds showed promising RhlR or LasR specific inhibitory activities over a range of concentrations below that causing toxicity. *ortho*-Vanillin, a previously untested compound, was the most promising within this set. Competition experiments against the native ligands for LasR and RhlR revealed that *ortho*-vanillin can interact competitively with RhlR but not with LasR. Overall, these studies expand our understanding of benzaldehyde activities in the LasR and RhlR receptors and reveal potentially promising effects of *ortho*-vanillin as a small molecule QS modulator against RhlR.

IMPORTANCE

Quorum sensing (QS) regulates many aspects of pathogenesis in bacteria and has attracted interest as a target for anti-virulence therapies. As QS is regulated by low molecular weight chemical signals, the development of chemical strategies that can interfere with this cell-cell communication pathway has seen considerable scrutiny over the past 25 years. Much of this research has focused on common human pathogens, including the LasR and RhlR QS receptors in *Pseudomonas aeruginosa*. Potent and selective chemical agents capable of blocking the activity of these receptors remain relatively scarce, however. Natural products have provided a bounty of chemical scaffolds with anti-QS activities, but their molecular mechanisms are poorly characterized. The current study serves to fill this void by examining the activity of an important and wide-spread class of natural product QS modulators, benzaldehydes and related derivatives, in LasR and RhlR. We demonstrate that *ortho*-vanillin can act as a competitive inhibitor of RhlR, a receptor that has emerged and may supplant LasR in certain settings as a target for QS control in *P. aeruginosa*. The results and insights provided herein will advance the design of chemical tools to study QS with improved activities and selectivities.

## INTRODUCTION

Many bacteria sense and respond to changes in population density using a gene regulation system called quorum sensing (QS). QS can regulate diverse behaviors including light production in marine bioluminescent bacteria, virulence factor production in plant and animal pathogens, and motility in many soil bacteria (1). In Proteobacteria, one type of QS system involves *N*-acyl L-homoserine lactone (AHL) signals (for reviews, see refs. (2, 3)). AHLs are produced by LuxI-type signal synthases, and detected by LuxR-type signal receptors, which are cytoplasmic transcriptional factors. At low population densities, AHLs are produced at low levels and accumulate in the local environment with increasing population density. The AHLs diffuse in and out of the cell, although active efflux can also contribute to the export of certain long chain AHLs (4). AHLs bind to the LuxR-type receptor protein and—for most of the known associative-type receptors—when they reach a critical concentration, they cause conformational changes to the protein that enable binding and activation of target gene promoters. AHLs interact with their cognate LuxR protein by making a series of hydrogen-bonding and hydrophobic contacts with residues in the ligand-binding pocket. AHL binding pockets vary structurally among LuxR family members to ensure specific responses to cognate AHLs, which differ in acyl chain structure.

*Pseudomonas aeruginosa* is an opportunistic pathogen that can cause debilitating infections in immunocompromised patients and is difficult to treat due to its multi-drug resistance. *P. aeruginosa* has two LuxI/R-type systems, LasI/R and RhlI/R. The LasI/R system produces and responds to *N-*(3-oxo)-dodecanoyl L-homoserine lactone (3OC12-HSL), and the RhlI/R system produces and responds to *N-*butanoyl L-homoserine lactone (C4-HSL). Upon AHL binding, LasR and RhlR activate distinct and overlapping regulons (5, 6). Among those are the genes encoding factors with known roles in virulence, such as the secreted toxins phenazine and hydrogen cyanide, proteases, and biofilm matrix proteins. These systems have been shown to be important for *P. aeruginosa* virulence in numerous acute animal infection models (7–11). Thus, *P. aeruginosa* QS has been proposed as an attractive target for the development of novel anti-virulence therapeutics (12).

Over the past 30 years, there has been considerable effort to identify molecules that block QS in *P. aeruginosa* and other bacteria. These prior studies have identified several promising approaches such as inhibiting LuxI-type synthases (13), destroying or sequestering AHLs (14), or inhibiting LuxR-type receptors (15). The latter strategy has received the most attention to date in *P. aeruginosa*, with much focus on the LasR receptor, and more recently RhlR, in *P. aeruginosa*. As a result, several promising molecules have been identified that inhibit these receptors (16–19). These molecules have potencies in the high-nM to mid- to low-μM range. In general, the most potent molecules have been identified as a result of high-throughput screens of small molecule libraries or by making targeted changes to the native AHL or other promising lead compounds via chemical synthesis.

In addition to these synthetic agents, there also has been widespread study of readily available molecules that can be re-purposed as QS inhibitors. Many of these compounds are natural products and were initially identified because of their ability to block QS-dependent phenotypes in the native species, not via studies of their ability to target specific QS pathways. These compounds include halogenated furanones (20), flavonoids such as baicalein (21, 22) and several benzaldehydes such as cinnamaldehyde (23–28). Despite the widespread use of these molecules as chemical tools for studies of QS inhibition, relatively little is known of the specificity, potency, and mechanism of action for most of these compounds. New tools to study

QS are of considerable interest, as many of the known chemical modulators have limitations, including relatively low potencies, efficacies, solubilities in aqueous media, and/or chemical stabilities. Consequently, re-purposed bioactive agents and readily-available natural products (and analogs) with promising QS inhibitory activities represents a valuable space to search for new chemical probes to study bacterial signaling.

In this study, we used *E. coli* reporters to evaluate the ability of several naturally occurring benzaldehydes and related derivatives to inhibit the *P. aeruginosa* QS receptors LasR and RhlR. We focused on compounds reported to disrupt QS-dependent phenotypes in *P. aeruginosa,* such as cinnamaldehyde and salicylic acid, along with several previously unstudied compounds with some structural similarity, such as orsellinaldehyde and *ortho*-vanillin (Fig. 1). We observed antagonism of the *E. coli* LasR and RhlR reporters at concentrations in the mid- to low-μM range, with *ortho*-vanillin showing the most promising effects. The compounds also caused mild reductions in growth and could nonspecifically antagonize a constitutive reporter at higher concentrations; however, at lower concentrations there was a suitable window of activity allowing for LasR and RhlR antagonism without any observable toxicity. In follow-up structure-function studies using LasR mutants, we found that critical AHL-binding residues in LasR were not required for *ortho*-vanillin to antagonize LasR. However, our results support that *ortho*-vanillin might specifically interact with RhlR. Together, our results indicate that naturally occurring benzaldehydes could have utility in QS inhibition and motivate future studies to develop this chemical scaffold into small-molecule tools to explore LuxR-type protein function and QS pathways.

**Fig. 1.**
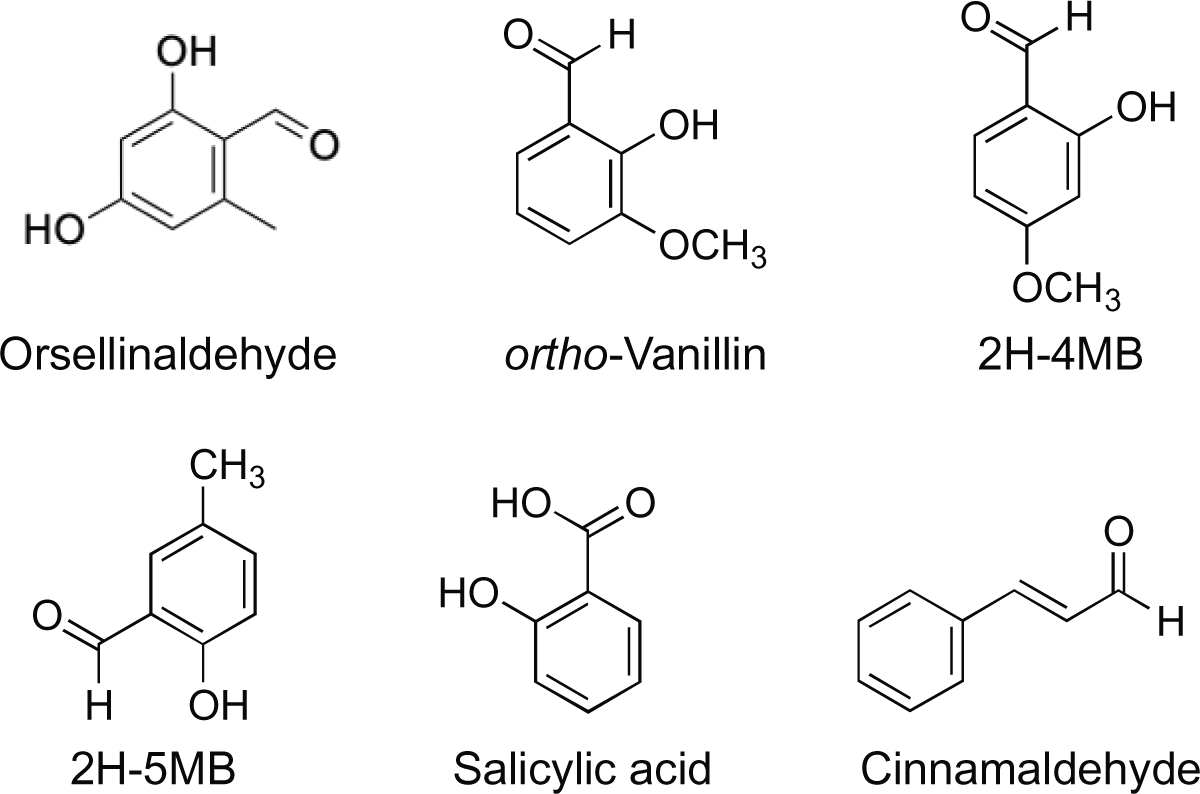
Structures of compounds examined in this study. 2H-4MB = 2-

## RESULTS

### Construction of cell-based *E. coli* bioreporters for LasR

To characterize compounds for their potential activity as LasR antagonists, we used a heterologous *Escherichia coli* strain expressing LasR from an arabinose-inducible promoter (P*ara-lasR)* on plasmid pJN105-L and a second plasmid with the LasR-inducible *lasI* promoter fused to a promoterless *lacZ* reporter (P*lasI-lacZ*) on plasmid pSC11-L (Fig. 2A). In this strain, *lacZ* expression required LasR and the LasR signal 3OC12-HSL (Fig. 2A), with a half-maximal activation concentration (i.e., EC_50_ value) of 65 nM. As a control, we also constructed an *E. coli* strain carrying a plasmid with *lacZ* expressed from the constitutive *aphA-3* promoter (29, 30), pVT19. With this strain, *lacZ* expression is fully activated in the absence of LasR or 3OC12-HSL (Fig. 2B).

**Fig. 2.**
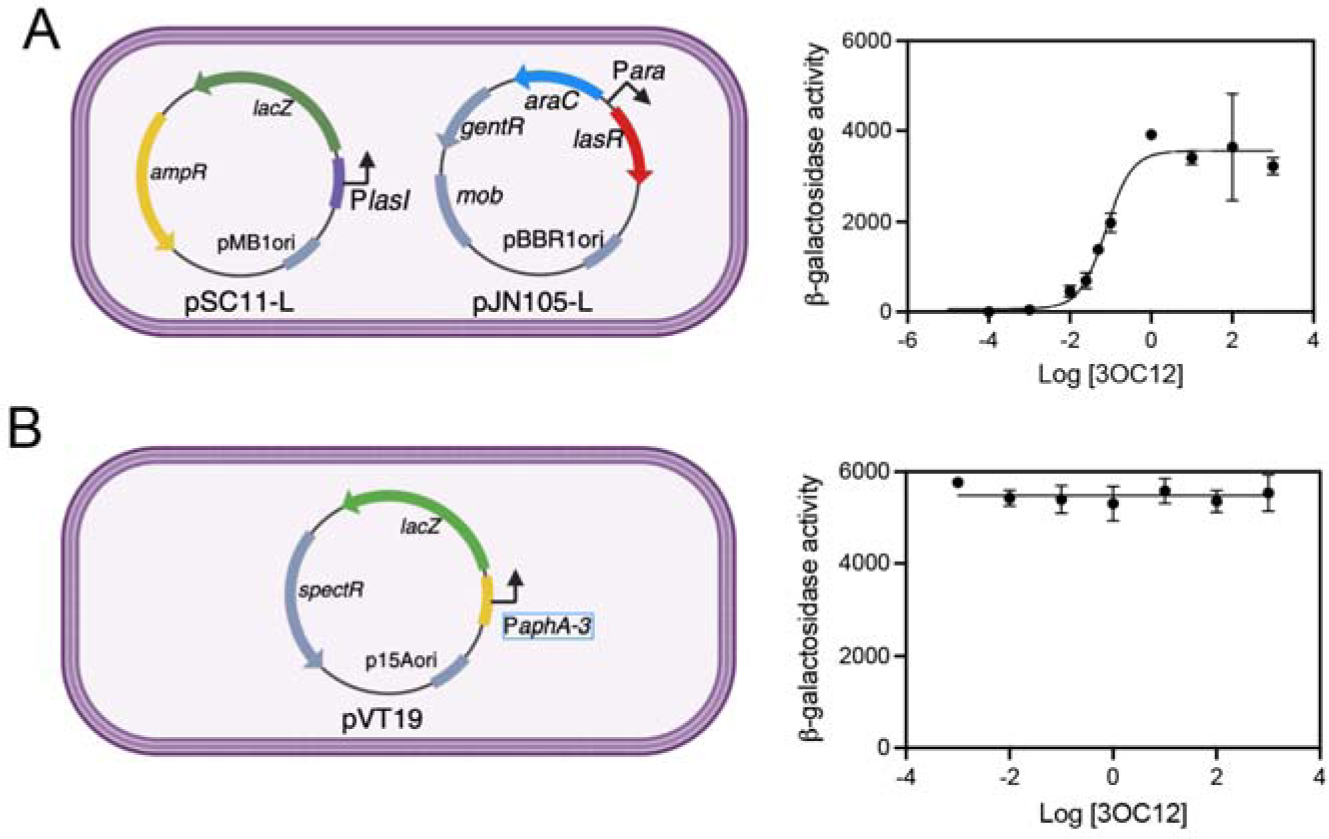
General schematic and initial characterization of *E. coli* bioreporter strains used for these studies. **A.** An *E. coli* bioreporter of LasR activity carries plasmid pSC11-L with a LasR-inducible P*lasI* promoter fused to the *lacZ* reporter and plasmid pJN105-L with an arabinose-inducible LasR. This strain produces β-galactosidase in response to 3OC12-HSL and 0.4% arabinose. The 3OC12-HSL induction is dose-responsive with a half-maximal concentration of 65 nM. **B.** An *E. coli* strain constitutively expressing *lacZ* carries the plasmid pVT19 with *lacZ* fused to the constitutive *aphA-3* promoter. This strain produces β-galactosidase in the presence or absence of 3OC12-HSL. Results are the averages of two (A) or three (B) independent experiments and the error bars represent the standard deviation.

### *E. coli* reporter assays indicate orsellinaldehyde antagonizes reporter activation nonspecifically

We utilized our *E. coli* reporters to evaluate the activity of the natural products and related derivatives in Fig. 1 as LasR antagonists, and we initiated our study with orsellinaldehyde, a metabolite produced by the fungus *Aspergillus nidulans* (31). Given its structural similarity to several known QS inhibitors, such as vanillin and salicylic acid, we were interested to examine orsellinaldehyde’s activity as a LasR antagonist. In the presence of 100 nM 3OC12-HSL (Fig. 3A and Table 1), we found that the concentration of orsellinaldehyde needed to inhibit P*lasI*-*lacZ* activity by 50% (i.e., its IC_50_ value) was 2374 µM (Fig. 3A, black line), indicating weak antagonist activity toward LasR. However, we observed that orsellinaldehyde caused a dose-dependent reduction of growth yield by about 10-20% at the highest concentrations (Fig. 3A, grey line). Further, orsellinaldehyde-dependent antagonism of the *lasI-lacZ* reporter correlated with its increasing effects on growth (correlation coefficient r=0.9877, p<0.0001, Fig. 3B). These results suggest antagonism of the LasR bioreporter by orsellinaldehyde may be due to the generalized effects of this compound on growth. To address this possibility, we generated a dose-response curve with orsellinaldehyde and our constitutive *lacZ-*producing control *E. coli* strain (with plasmid pVT19). We found orsellinaldehyde antagonized the constitutive *lacZ* reporter in a dose-responsive manner with an IC_50_ of 2308 µM (Fig. 3A, red line), which was similar to that of the LasR bioreporter (2374 µM). These results support the conclusion that orsellinaldehyde antagonizes *lacZ* reporter activation in a nonspecific manner.

**Fig. 3.**
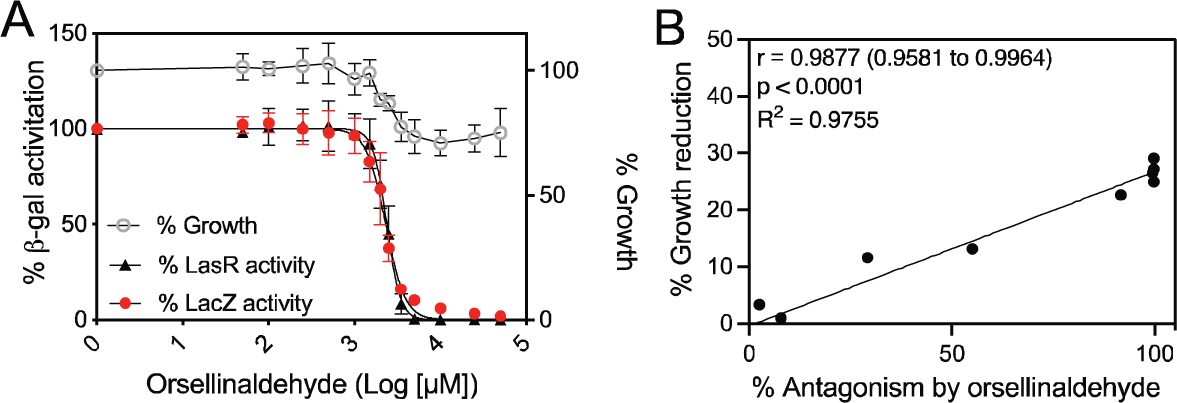
Activity of orsellinaldehyde with *E. coli* LasR and constitutive reporters. **A**. Grey line, orsellinaldehyde-dependent growth inhibition as a percent of cells with no orsellinaldehyde. Dose-response curve of orsellinaldehyde in competition with 100 nM 3OC12-HSL in cultures of *E. coli* with arabinose-inducible LasR and a LasR-dependent *lasI-lacZ* reporter (black line) or of *E. coli* with a constitutive *aphA-3-lacZ* reporter (red line). IC50 values are given in Table 1. Results show the averages of five (LasR) or three (LacZ control) independent experiments and the error bars represent the standard deviation. **B.** Average values from A (% growth reduction vs. % inhibition of the *lasI-lacZ* reporter), which were used to determine Pearson’s correlation coefficient (r value) and significance (p) and generate a fitted line using a simple linear regression model.

**Table 1.**
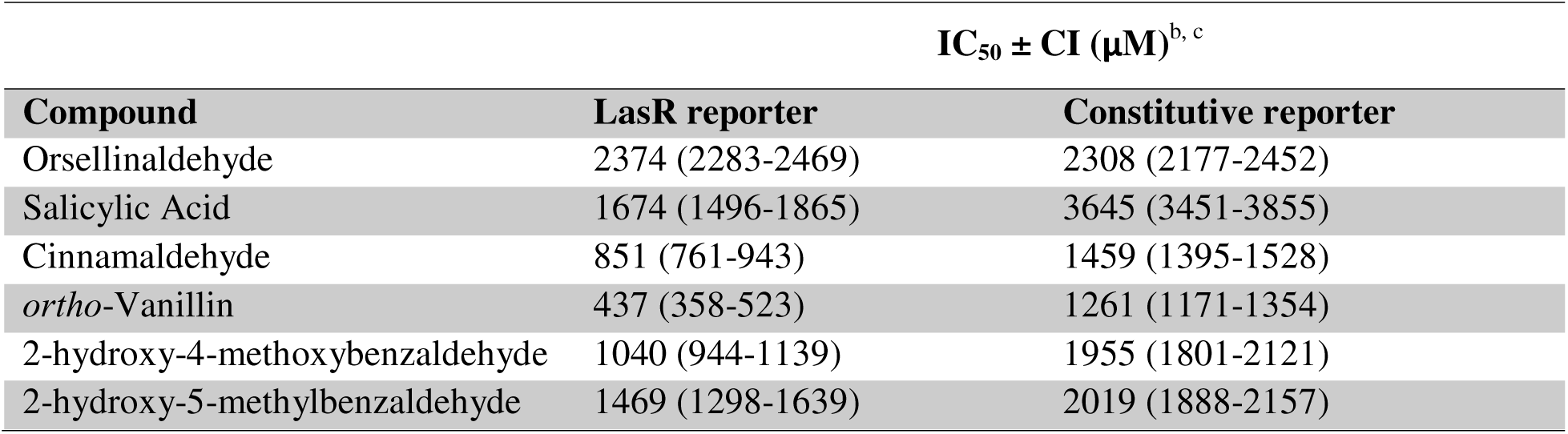
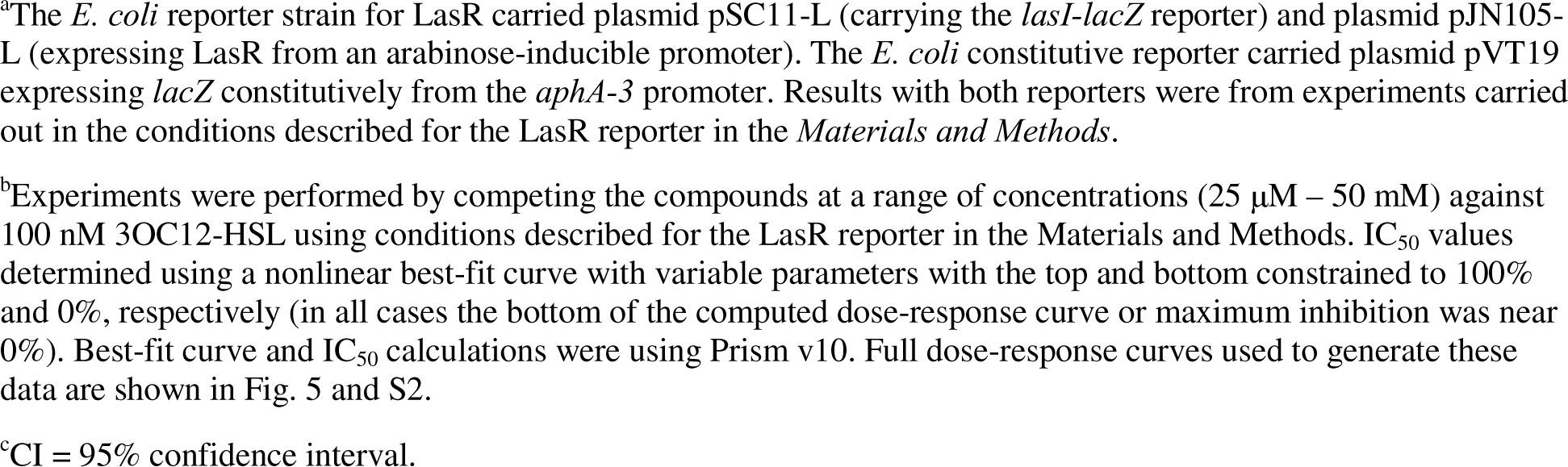
Potency of benzaldehydes using *E. coli* LasR and constitutive reporters^a^.

| Compound | IC<sub>50</sub> ± CI (μM)<sup>b, c</sup> |  |
| --- | --- | --- |
|  | LasR reporter | Constitutive reporter |
| Orsellinaldehyde | 2374 (2283-2469) | 2308 (2177-2452) |
| Salicylic Acid | 1674 (1496-1865) | 3645 (3451-3855) |
| Cinnamaldehyde | 851 (761-943) | 1459 (1395-1528) |
| ortho-Vanillin | 437 (358-523) | 1261 (1171-1354) |
| 2-hydroxy-4-methoxybenzaldehyde | 1040 (944-1139) | 1955 (1801-2121) |
| 2-hydroxy-5-methylbenzaldehyde | 1469 (1298-1639) | 2019 (1888-2157) |

<sup>a</sup>The *E. coli* reporter strain for LasR carried plasmid pSC11-L (carrying the *lasI-lacZ* reporter) and plasmid pJN105-L (expressing LasR from an arabinose-inducible promoter). The *E. coli* constitutive reporter carried plasmid pVT19 expressing *lacZ* constitutively from the *aphA-3* promoter. Results with both reporters were from experiments carried out in the conditions described for the LasR reporter in the *Materials and Methods*.
<sup>b</sup>Experiments were performed by competing the compounds at a range of concentrations (25 μM – 50 mM) against 100 nM 3OC12-HSL using conditions described for the LasR reporter in the Materials and Methods. IC<sub>50</sub> values determined using a nonlinear best-fit curve with variable parameters with the top and bottom constrained to 100% and 0%, respectively (in all cases the bottom of the computed dose-response curve or maximum inhibition was near 0%). Best-fit curve and IC<sub>50</sub> calculations were using Prism v10. Full dose-response curves used to generate these data are shown in Fig. 5 and S2.
<sup>c</sup>CI = 95% confidence interval.

To test whether these effects were specific to the *lacZ* reporter or general to other reporters, we generated an orsellinaldehyde dose-response curve using a strain constitutively expressing GFP (*E. coli* carrying a constitutive GFP-producing plasmid pUC18T-mini-Tn7T-Gm-gfpmut3). Orsellinaldehyde also antagonized the constitutive GFP reporter with an IC_50_ of 1057 µM for GFP (Fig. S1), which was similar to that of the *lacZ* reporter. These results support the conclusion that the effects of orsellenaldehyde on our LasR bioreporter are related to a generalized effect on gene expression or other cellular processes and not specific to LasR.

### Evaluation of other benzaldehyde derivatives in *E. coli* LasR reporters

We next examined compounds structurally related to orsellinaldehyde and previously reported to modulate QS for antagonistic activity in LasR. In view of the results above, we questioned whether some of the reported inhibitory activities were also largely due to nonspecific toxic effects. We selected several such compounds; cinnamaldehyde (25), salicylic acid (25–28), and the as-yet untested, but related compounds *ortho*-vanillin, 2-hydroxy-5-methylbenzaldehyde and 2-hydroxy-4-methoxybenzaldehyde (Fig. 1). The results (Fig. 4, Fig. S2 and Table 1) show that each of the compounds can antagonize the LasR-dependent *lasI-lacZ* reporter with IC_50_s ranging from 437 µM for vanillin to 1674 µM for salicylic acid. We also observed decreases in growth like that of orsellinaldehyde by ∼25% at the highest concentrations (Fig. 4). The effects on growth and inhibition of the *lasI-lacZ* reporter were significantly correlated for each of the compounds (Fig. 4, Fig. S2 and Table 1), although there was a weaker correlation for *ortho*-vanillin and cinnamaldehyde because the effects on growth were minimal at the lower concentrations (Fig. 4A and C, right side). We also generated dose-response curves of each compound with the control constitutive *lacZ* reporter strain (Fig. 4). All of these compounds were less potent with the constitutive reporter than that of the LasR reporter, by 1.3-fold for 2-hydroxy-5-methylbenzaldehyde to almost 3-fold lower for *ortho*-vanillin. These results suggest that, while all of the compounds also have nonspecific effects at higher concentrations, certain compounds—i.e., *ortho*-vanillin and cinnamaldehyde—have some specific activity against LasR at lower concentrations.

**Fig. 4.**
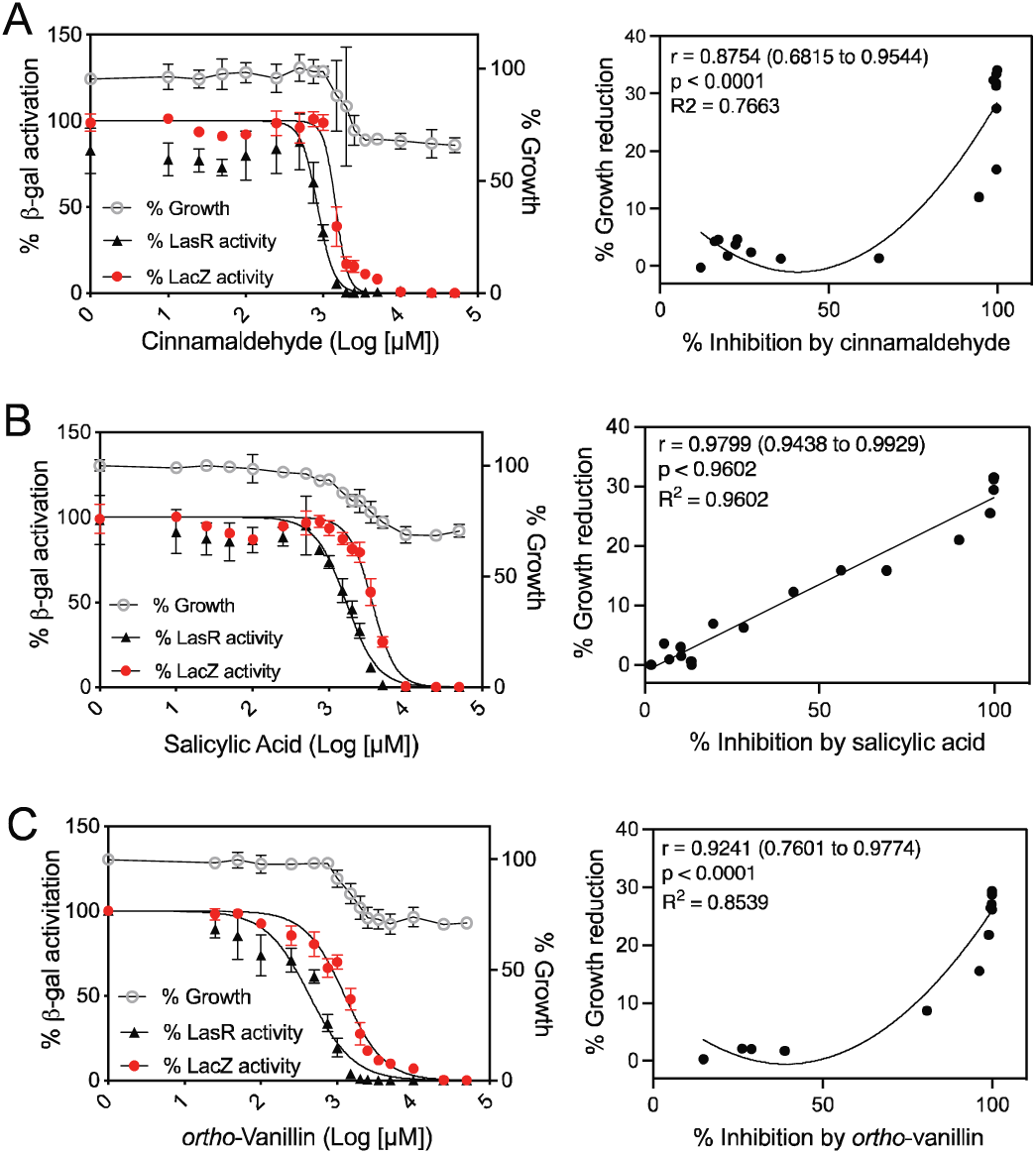
Activities of cinnamaldehyde (A), salicylic acid (B) and vanillin (C) with *E. coli* LasR and constitutive reporters. Left column: dose-response curves for each indicated compound in competition with 100 nM 3OC12-HSL in cultures of *E. coli* with arabinose-inducible LasR and a LasR-dependent *lasI-lacZ* reporter (black line) or of *E. coli* with a constitutive *aph-lacZ* reporter (red line). Results show the averages of four independent experiments and the error bars represent the standard deviation. IC_50_ values from the fit curves are given in Table 1. Right column shows average values from the graphs on the left (% growth reduction vs. % inhibition of the *lasI-lacZ* reporter), which were used to determine Pearson’s correlation coefficient (r value) and significance (p) and generate a fitted line using a simple linear regression model (B) or a second-order polynomial nonlinear regression model (A and C).

### Results of LasR mutant reporters support *ortho*-vanillin not contacting specific residues in the LasR ligand-binding domain

As *ortho*-vanillin was the most potent LasR antagonist identified above, we sought to further characterize the nature of potential *ortho*-vanillin/LasR interactions. To our knowledge, no other studies have experimentally addressed the molecular mechanism by which benzaldehyde derivatives antagonize LuxR-type receptors. We began by asking whether *ortho*-vanillin is acting as a competitive LasR antagonist, similar to the synthetic compound V-06-018 (19), and binding in the native ligand (i.e., 3OC12-HSL) binding site. To this end, we applied an approach of competing *ortho*-vanillin with 3OC12-HSL at varying concentrations using our LasR reporter assay described above. The reporter experiments in the heterologous *E. coli* host provide a proxy to assess LasR interaction with AHLs in the absence of other host regulation effects. The ability of *ortho*-vanillin to antagonize LasR should vary when the 3OC12-HSL concentration is increased if both molecules are competing for binding to the same site in LasR. We generated antagonism dose-response curves for *ortho-*vanillin competed against 3OC12-HSL at 100 nM, 10 μM and 100 μM (Fig. 5A). Although there was a small difference in the *ortho-* vanillin IC_50_ at 100 nM and 10 μM, this difference was not significant (p>0.07). These results do not support the conclusion that *ortho-*vanillin is a competitive antagonist of LasR.

**Fig. 5.**
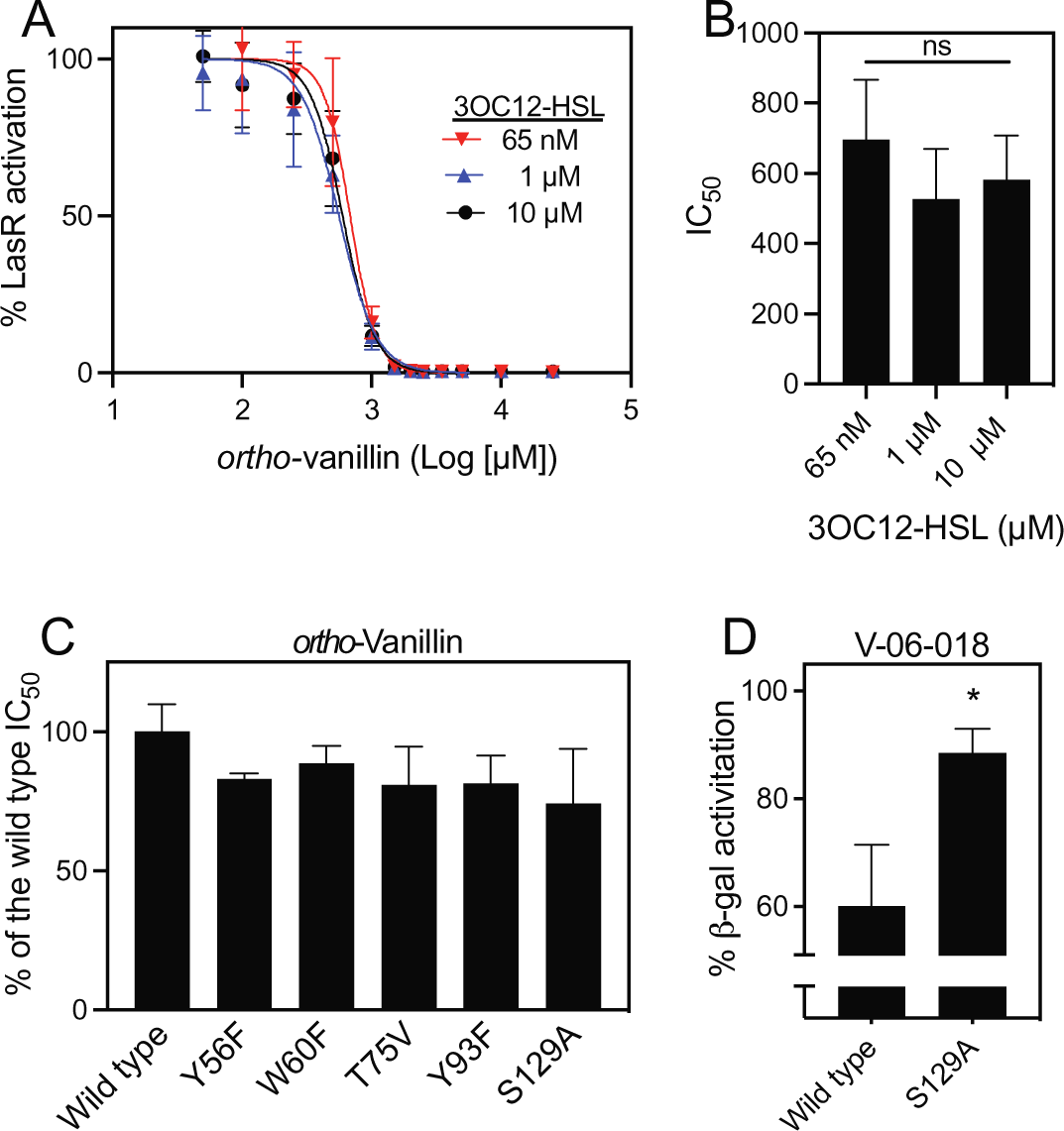
LasR mutant antagonism data for *ortho-*vanillin and V-06-018. **A.** Dose-response curves of *o-* vanillin in competition with 3OC12-HSL at its EC_50_ value of 65 nM, 1 μM or 10 μM in *E. coli* with arabinose-inducible LasR and a LasR-dependent *lasI-lacZ* reporter. Each curve shows results of three independent experiments with the standard deviation represented by horizontal bars. **B.** Data show averages of IC_50_ values from each curve shown in panel A. Error bars show the standard deviation. There were no statistical differences between any of the conditions by one-way ANOVA (p>0.3). **C.** *ortho-* Vanillin was tested at varying concentrations (25 nM to 50 μM) against 3OC12-HSL at its EC_50_ value in the specific *E. coli* reporter strain as indicated on the X-axis. The *ortho-*vanillin dose-response curves (shown in Fig. S6) were used to determine the IC_50_ values, which are shown as the % of the IC_50_ of *ortho*-vanillin for *E. coli* with wild type LasR. There were no significant differences of any of the LasR mutant results from that of wild type by one-way ANOVA. **D.** The LasR inhibitor V-06-018 was tested at 100 μM against 3OC12-HSL at its EC_50_ value (65 nM) in *E. coli* with wild type or the S129A LasR as indicated on the X-axis. Values are reported as the % of reporter activation with the EC_50_ of 3OC12-HSL with no other compound. *, statistical significance by students t-test (p<0.05).

In addition, we performed *in silico* docking studies of *ortho-*vanillin within the ligand-binding site (LBD) of LasR using the reported full-length LasR structure (PDB ID: 6V7X; see Methods) and found that this compound could be accommodated. Three residues were identified that could be important for the *ortho-*vanillin/LasR interaction: Thr75, Thr115 and Ser129 (Fig. S3). These residues were predicted to hydrogen bond with the phenol and aldehyde substituents of *ortho*-vanillin. Several other residues, such as Tyr56, Trp60, and Tyr93 were also predicted to form close contacts with *ortho*-vanillin. Ser129 and several other of these residues (e.g., Arg61, Tyr56 and Asp73) were also found to be important for LasR interaction with 3OC12-HSL and other ligands (19, 32, 33) (Fig. S3).

To examine these putative interactions between *ortho-*vanillin and LasR, we determined the activity of *ortho-*vanillin in several LasR mutants. In prior studies in our laboratories, a set of LasR mutants were generated in which residues within the ligand-binding pocket were mutated to a different residue of similar steric size but without the capability to hydrogen bond (e.g., Tyr ➔ Phe). These mutants were introduced into *E. coli* to generate *lasI-lacZ* reporters analogous to the wild-type LasR reporter above (see Table S1 and Methods). From that set, we selected five LasR mutant reporters to test *ortho*-vanillin (W60F, Y56F, T75V, Y93F, and S129A), which included the Thr75 and Ser129 residues predicted to be important for *o-*vanillin interaction in our *in silico* study. Each of these mutants showed varying degrees of activation by 3OC12-HSL in our reporter experiment (Fig. S4), which was consistent with prior results (33). We generated dose-response curves with *ortho-*vanillin competed against 3OC12-HSL at the concentration needed to cause half-maximal LasR activation (EC_50_) for each mutant (Fig. S4). In our experiments, *ortho-*vanillin antagonism of the LasR mutants was indistinguishable from that of the wild type LasR (Fig. 5C and S6). As a control, we also tested the ability of V-06-018 to antagonize the LasR S129A mutant. Consistent with prior results (19), we showed that V-06-018 was significantly less potent with the S129A mutant compared with wild type LasR (Fig. 5D).

Together, these results show that some of the LasR ligand binding site residues that make important contacts with other ligands (AHL and non-AHL agonists or antagonists) are not required for *ortho-*vanillin activity and are consistent with the idea that *ortho-*vanillin does not interact with the LasR ligand-binding domain in a mode analogous to other ligands.

### Evaluation of benzaldehyde derivatives using *E. coli* RhlR reporters indicate vanillin can antagonize RhlR

We hypothesized that the benzaldehyde derivatives in our studies (Fig. 1) could be poor antagonists of LasR because they have very short or no acyl tail functionality, which has been shown to be important for LasR interactions in studies of the native ligand 3OC12-HSL and other inhibitors, such as V-06-018 (19, 34). We thus turned our attention to RhlR from *P. aeruginosa*, which is regulated by an AHL with a much shorter 4-carbon tail, C4-HSL. We performed *in silico* docking studies analogous to those for LasR above using the recently published RhlR structure, which was purified with a non-native agonist meta-bromothiolactone (mBTL) (PDB ID: 8DQ0) (35). We examined docking of *ortho-*vanillin and the native ligand C4-HSL to RhlR (Fig. S6), and found that both could be accommodated. The phenol moiety of *ortho-*vanillin was predicted to hydrogen bond with Asp81 of RhlR, supporting the idea that this compound could possibly interact with RhlR. The docking score calculations were similar for *ortho-*vanillin and the native ligand C4-HSL (about -5.2 kcal/mol), although the specific interactions with RhlR appeared to be different for C4-HSL, which was predicted to have close contact with residues Tyr69, Trp93 and Ala108, but not with Asp81.

To test the ability of our set of benzaldehydes and related compounds to antagonize RhlR, we generated dose-response curves with these compounds using an *E. coli* RhlR reporter strain (Fig. 6 and Table 2). This strain is analogous to the LasR reporter above but it carries plasmid pECP61.5 expressing RhlR from the IPTG-inducible P*lac* promoter as well as the *rhlA-lacZ* reporter (36). We also utilized our constitutive *lacZ* reporter plasmid pVT19 to generate dose-response curves using the RhlR assay conditions (Fig. 6 and Table 2). The tested compounds caused maximal ∼20% growth reduction for the RhlR conditions. The potencies of our compounds with the RhlR reporter ranged from an IC_50_ of 151 μM for *ortho-*vanillin to ∼10 mM for salicylic acid. With the constitutive *lacZ* reporter, there was a similar spread in potencies for our compounds, with *ortho-*vanillin having the lowest IC_50_ and salicylic acid as the highest.

**Fig. 6.**
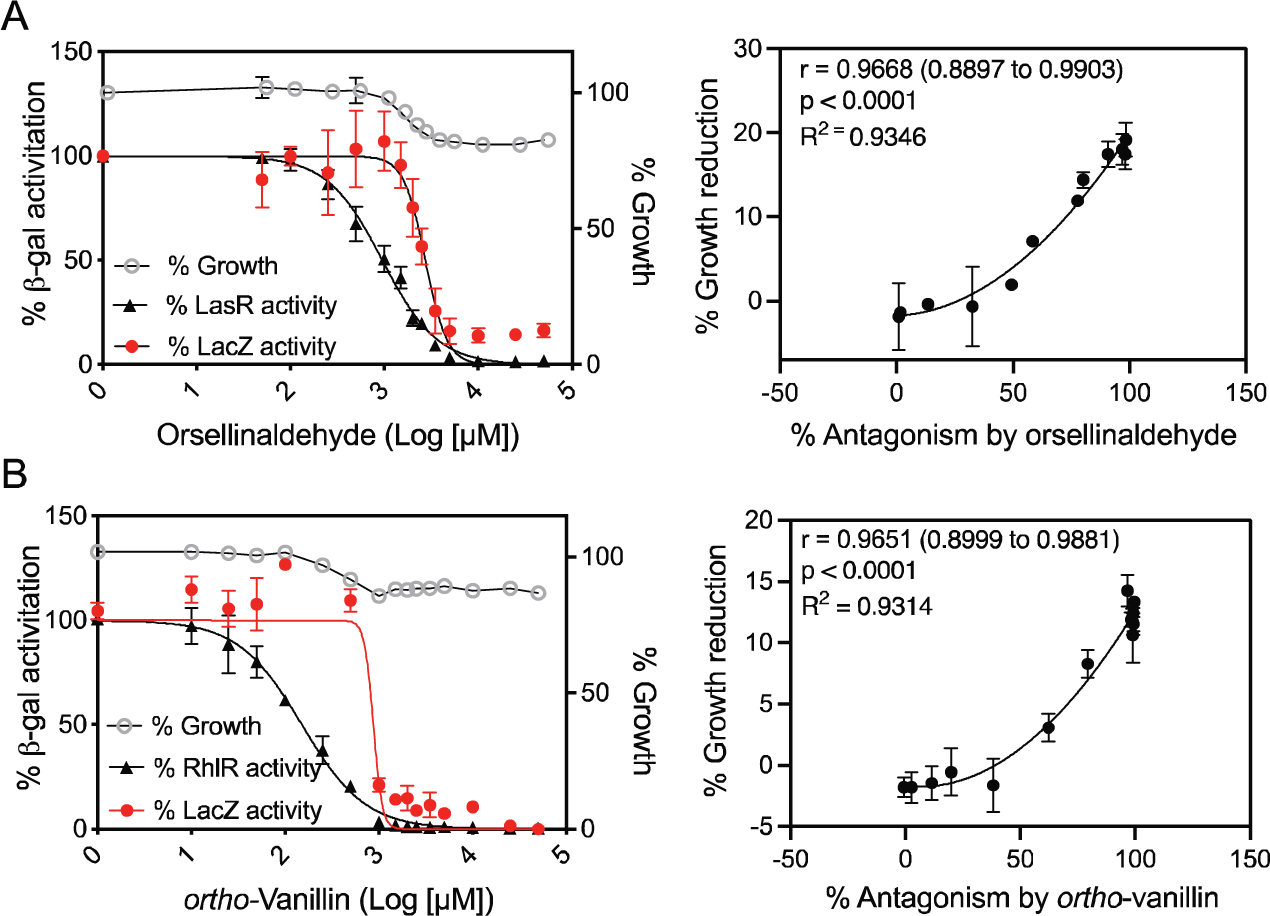
Activities of orsellinaldehyde (A) and *o-*vanillin (B) with *E. coli* RhlR and constitutive reporters. Left column: dose-response curves for each indicated compound in competition with 414 nM C4-HSL in cultures of *E. coli* with IPTG-inducible RhlR and a RhlR-dependent *rhlA-lacZ* reporter (black symbols) or of *E. coli* with a constitutive *aph-lacZ* reporter (red symbols). Results show the averages of three independent experiments and the error bars represent the standard deviation. IC_50_ values from the fit curves are given in Table 2. Right column shows average values from the graphs on the left (% growth reduction vs. % *rhlA-lacZ* reporter antagonism), which were used to determine Pearson’s correlation coefficient (r value) and significance (p) and generate fitted lines using a second-order polynomial nonlinear regression model.

**Table 2.**
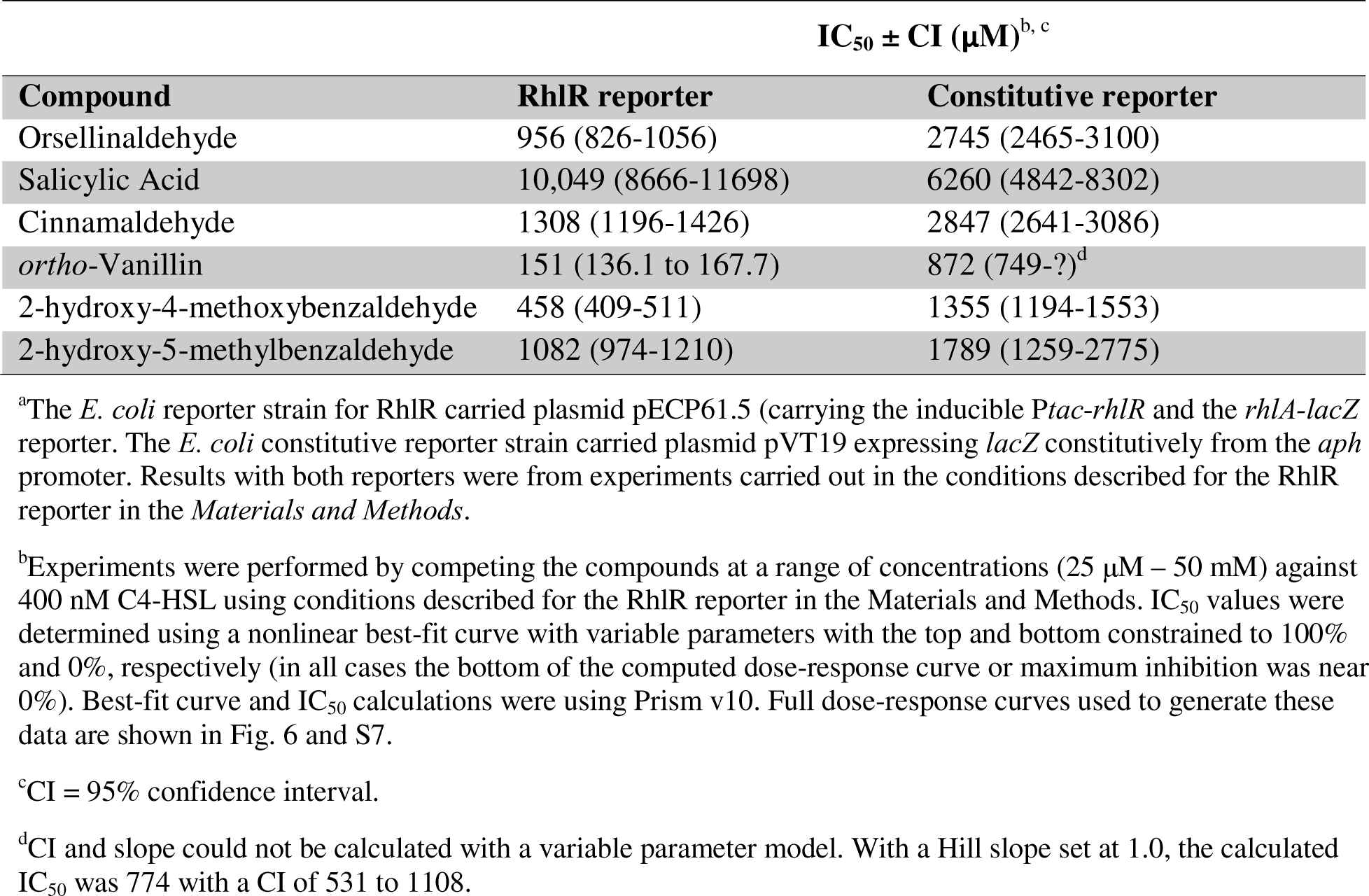
Potency of benzaldehyde derivatives using *E. coli* RhlR and constitutive reporters^a^.

However, the IC_50_ for *ortho-*vanillin was 5-fold lower with RhlR than with the constitutive *lacZ* reporter. There were also no observed effects of *ortho-*vanillin on growth until concentrations at which there was >50% antagonism of the RhlR reporter, and there was no antagonism of the constitutive *lacZ* reporter until concentrations >500 μM. These results support the idea that *ortho-*vanillin may specifically antagonize RhlR at concentrations below 500 μM.

### RhlR reporter data support a competitive mechanism of RhlR antagonism by *ortho-*vanillin

We were interested to determine whether *o-*vanillin was acting as a competitive RhlR antagonist. As with LasR, we tested whether competing with C4-HSL at different concentrations could elicit changes in the ability of *ortho-*vanillin to antagonize RhlR in the *E. coli lacZ* reporter. We generated antagonism dose-response curves for *ortho-*vanillin competed against C4-HSL at 400 nM, 10 μM, and 100 μM (Fig. 7). We observed a significant C4-HSL concentration-dependent decrease in the potency of *ortho*-vanillin. These differences were most apparent at the lowest concentrations of *ortho*-vanillin, which were below the concentration at which nonspecific antagonism of the *lacZ* reporter were observed. These results are congruent with the ability of *ortho*-vanillin can act as a competitive antagonist of RhlR.

**Fig. 7.**
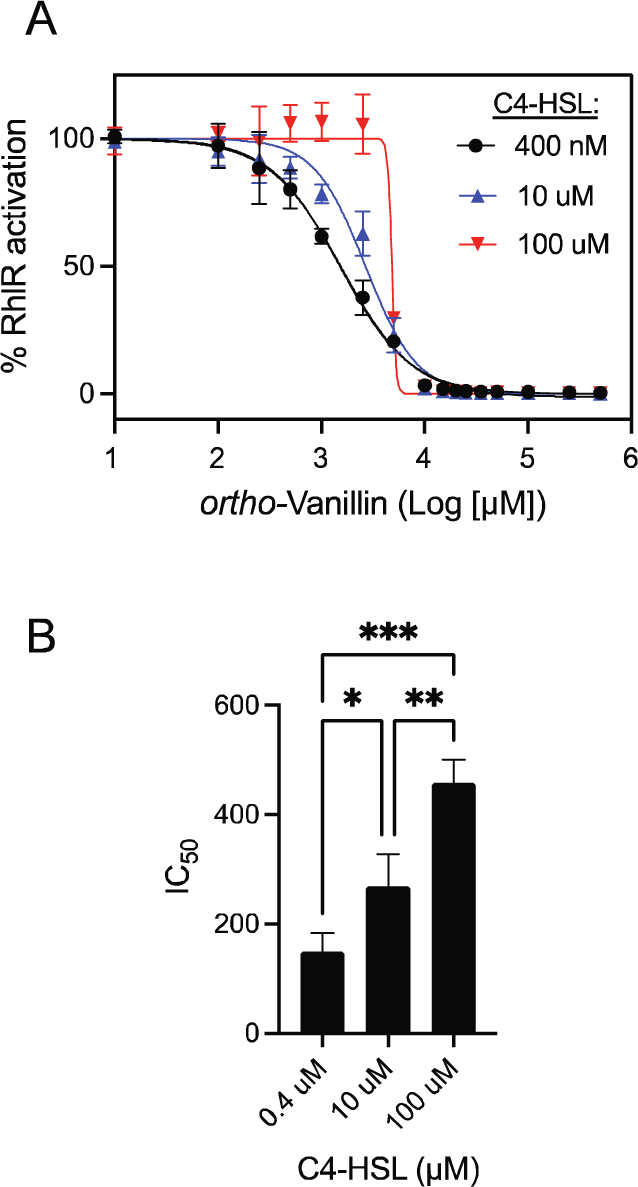
Dose-response RhlR antagonism by *ortho-*vanillin competed with varying C4-HSL concentrations. **A.** Dose-response curves of *ortho-*vanillin in competition with 400 nM, 10 μM or 100 μM C4-HSL in cultures of *E. coli* with IPTG-inducible RhlR and a RhlR-dependent *rhlA-lacZ* reporter. Each curve shows results of three independent experiments with the standard deviation represented by horizontal bars. **B.** Data show averages of IC_50_ values from each curve shown in panel A. Error bars show the standard deviation. Statistical significance by one-way anova; *, p<0.05; ** p<0.01; *** p<0.001.

## DISCUSSION

The contribution of QS to a wide array of phenotypes, including virulence, in *P. aeruginosa* has attracted significant attention to the identification of QS inhibitors for use as chemical probes and in therapeutic development. Despite considerable work in this area, there are relatively few highly potent and selective QS inhibitors in *P. aeruginosa* and related proteobacteria. Most of these compounds target LuxR-type receptor proteins, including V-06-018 that antagonizes LasR in *P. aeruginosa* (18) and the chlorolactone AHL analog (CL) that antagonizes CviR from *Chromobacterium violaceum* (37, 38). Beyond these classes of synthetic compounds, there are many naturally derived compounds or extracts that have reported activities as QS inhibitors in bacteria. For example, salicylic acid can downregulate production of the QS-controlled virulence factors pyocyanin and elastase and attenuate the ability of *P. aeruginosa* to infect plants (28). However, detailed studies to determine the molecular mechanisms by which these natural products elicit their effects on QS are limited. In this study, we evaluate the ability of salicylic acid, cinnamaldehyde, and several related benzaldehyde derivatives, to antagonize the *P. aeruginosa* LuxR-type receptors LasR and RhlR using heterologous reporters in *E. coli*.

We provide evidence that one of these compounds, namely *ortho-*vanillin, can specifically antagonize these receptors within a lower range of concentrations in which they are not generally toxic. These results provide a basis to guide the use of these compounds in QS studies and suggest chemical scaffolds to advance for the design of new QS receptor antagonists.

The investigations described here indicate that *ortho-*vanillin can specifically antagonize LasR and that it does so through a non-competitive mechanism (Fig. 5). There are prior reports of other compounds that might inhibit LuxR-type receptors noncompetitively. Halogenated furanones, such as bromofuranone, have been shown to inhibit the *Vibrio fischeri* LuxR receptor noncompetitively (39). Inhibition might involve a mechanism of increasing turnover of the receptor protein in the cell (39), although bromofuranone can also be broadly toxic at inhibitory concentrations (40). Some flavonoids also have been reported to inhibit LasR noncompetitively, such as baicalein, although in the case of baicalein the mechanism is not known (22). Our discovery that *ortho*-vanillin can antagonize LasR noncompetitively adds to this list of noncompetitive antagonists.

In the case of RhlR, *ortho*-vanillin appears to act as a specific, competitive antagonist in the *E. coli* reporter (Fig. 7). Competitive inhibition is by far the most invoked mechanism for known LuxR-type inhibitors; the crystal structure of chlorolactone (CL) bound to CviR and stabilizing an inactive conformation provides perhaps the most compelling support for this mechanism (38). There are several other known competitive inhibitors of RhlR, most of which closely resemble its native ligand C4-HSL, and our prior detailed structure-function studies have revealed portions of the molecules that are essential for strong inhibitory activity (41). With the recently determined crystal structure of RhlR (35), it is now possible to carry out more detailed studies to better understand RhlR-ligand binding interactions, including with the native ligand C4-HSL. Such studies will be interesting to reveal important insight into the mechanism of RhlR-ligand interactions and advance the design of compounds that can modulate RhlR activity.

Our results with *E. coli* reporters show that *ortho-*vanillin is more potent against RhlR than LasR. This difference could be due to the relatively small size of this molecule and/or its lack of an acyl tail. The natural ligand of LasR, 3OC12-HSL, has a long 12-carbon acyl tail, whereas the RhlR ligand C4-HSL has a much shorter 4-carbon acyl tail. Prior structure-function studies of LasR and 3OC12-HSL reveal that there are important hydrophobic contacts formed between the long tail of 3OC12-HSL and residues within the LasR binding pocket (42). These contacts contribute to the strength and specificity of the interaction with LasR. In addition, studies with V-06-018 analogs showed that shorter acyl tails weaken LasR interactions (43). In turn, we have shown that RhlR is both activated and inhibited by AHLs analogs with shorter tails. *ortho-*Vanillin largely lacks such a hydrophobic tail (Fig. 1), which might weaken its ability to antagonize LasR, while enhance its ability to engage with RhlR. Our results support the idea that the hydrophobic tails of ligands play a critical role in the specificity and strength of interactions with LuxR proteins. As this competitive activity for *ortho*-vanillin in RhlR, and its non-competitive activity in LasR, were observed in *E. coli* reporter systems, additional experiments including *in vitro* studies will be necessary to provide further clarity into its molecular mechanisms of action and the hypotheses outlined here. The relative simplicity of the *ortho*-vanillin scaffold suggest straightforward routes to alter its structure and examine impact on potency and specificity, along with reducing any associated toxicity. Overall, these studies illustrate the importance of performing rigorous studies to determine the specificity and function of small molecule QS inhibitors to inform their use as research tools and other applications.

## MATERIALS AND METHODS

### Culture conditions and reagents

Unless otherwise noted, bacteria were grown at 37 °C in Lysogeny broth (LB; 10 g tryptone, 5 g tryptone and 5 g NaCl per L), or on LB agar (LBA; 1.5% (weight per volume) Bacto-Agar). For RhlR bioreporter experiments, growth was at 30 °C and in A medium (44)(60 mM K_2_HPO_4_, 33 mM KH_2_PO_4_, 7.5 mM (NH_4_)2SO_4_, 1.7 mM sodium citrate ·2H_2_O, 0.4% glucose, 0.05% yeast extract, 1 mM MgSO_4_). All *E. coli* broth cultures were grown with shaking at 250 rpm, 18 mm test tubes (for 5 ml cultures) or 125 ml baffled flasks (for 10 ml cultures) unless otherwise specified. For selection, 100 µg ml^-1^ ampicillin, 10 µg ml^-1^ gentamicin, or 150 µg ml spectinomycin were used. For experiments with the RhlR bioreporter strain, A medium was used as described (36, 45). When needed for induction of LasR or RhlR, we added IPTG (isopropyl β-D-1-thiogalactopyranoside) at 1 µM final concentration and L-(+)-arabinose at 0.25% final concentration. Native HSLs were suspended in ethyl acetate acidified with 0.01% glacial acetic acid and added to culture tubes and dried down prior to adding growth medium for experiments.

We measured β-galactosidase activity with a Tropix Galacto-Light Plus chemiluminescence kit according to the manufacturer’s protocol (Applied Biosystems, Foster City, CA). Native HSLs (3oxoC12-HSL and C4-HSL) were purchased from Cayman Chemicals (MI, USA), gentamicin was purchased from GoldBio (MO, USA) and ampicillin and spectinomycin were purchased from Sigma Aldrich (MO, USA). DMSO (solvent for inhibitor compounds), IPTG, and L-(+)-arabinose were purchased from Fisher Scientific (PA, USA). Natural products and benzaldehyde derivatives were purchased from Sigma Aldrich (MO, USA). V-06-018 was synthesized as previously described (19).

### Strains and plasmids

Strains and plasmids are listed in Table S1. To assess LasR activation of *lasR* expression in recombinant *E. coli*, we used *E. coli* strain DH5α carrying two plasmids; plasmid pJN105-L (46) with an arabinose-inducible *P. aeruginosa lasR* and plasmid pSC11-L (47) with the promoter of the LasR-responsive gene *lasI* fused to a *lacZ* reporter. For some studies, pJN105-L was replaced with derivatives of this plasmid encoding LasR mutants with single amino acid substitutions (32). To assess RhlR activation of *rhlR* expression in recombinant *E. coli*, we used *E. coli* DH5α with plasmid pECP61.5 (36) with an IPTG-inducible p*tac-rhlR* and a RhlR-responsive gene *rhlA* fused to the *lacZ* reporter. For constitutive expression of the *lacZ* reporter, we used *E. coli* DH5α with plasmid pVT19, which has the *lacZ* gene fused to the constitutive *aphA-3* promoter. To construct pVT19, the constitutive *aphA-3* promoter (29, 30) was amplified from a pTCV-lac derivative using primers Vlac1 and Vlac2 (30). The resulting amplicon was digested with EcoRI and BamHI and ligated into similarly digested pKS12A (48). The resulting plasmid with the *aphA-3* promoter transcriptionally fused to *lacZ* was designated pVT19. For constitutive expression of the *gfp* reporter, we used *E. coli* DH5α with plasmid pUC18T-mini-Tn7T-Gm-gfpmut3 (49).

### Transcription reporter assays in *E. coli*

To assess LasR activation of *lasR* expression in recombinant *E. coli*, overnight cultures of *E. coli* DH5α pSC11-L, pJN105-L were diluted 1:100 into LB containing selection antibiotics gentamicin and ampicillin in 10 ml cultures. When the cultures reached an OD_600_ of 0.2-0.3, L-(+)-arabinose was added to a final concentration of 0.25%. The control did not receive L-(+)-arabinose. The cultures were then grown to an OD_600_ of 0.5-0.6 and 500 µL was added to 1.5 mL micro centrifuge tubes containing dried 3OC12-HSL. Aliquots (5 µL) of increasing concentrations of inhibitor test compound stock solution in DMSO were then added to the designated micro centrifuge tubes containing culture. Tubes containing *E. coli* with just signal and DMSO were included as controls. After 3 h at 37 °C with shaking, OD_600_ was measured using a plate reader and β-galactosidase activity was measured as described above.

To assess RhlR activation of *rhlR* expression in recombinant *E. coli*, overnight cultures of *E. coli* DH5α pECP61.5 grown at 30 °C in A medium containing antibiotic selection (ampicillin) and IPTG to induce RhlR expression were diluted to an OD_600_ of 0.1 and 1 mL was added to culture tubes containing dried C4-HSL. Aliquots (5 µL) of DMSO containing increasing concentrations of inhibitor test compound or DMSO with no test compound were added to the designated Eppendorf tubes containing culture. Tubes containing *E. coli* with signal and DMSO were included as a vehicle control. After 5 h at 30 °C with shaking, OD_600_ was measured using a plate reader and β-galactosidase activity was measured as described above.

Experiments with the LasR mutants and the constitutive *lacZ* expression plasmid pVT19 or constitutive *gfp* expression plasmid pUC18T-mini-Tn7T-Gm-gfpmut3 were carried out identically as described above for the LasR or RhlR bioreporter experiments. Results with the pBT19 constitutive reporter strain was different for the LasR vs. RhlR bioreporter protocols likely due to differences in growth conditions (temperature and/or growth media).

### Computational modeling

The structure of LasR (PDB ID: 6V7X) and RhlR (PDB ID: 8DQ0) was used for docking studies using the Lamarckian protocol and the empirical free energy function in AutoDock version 4.2. The hit search was refined using an improved docking method. The α-β-α sandwich located near the N-terminal ligand binding domain (LBD) was used as the binding location for docking calculations. The protein target was prepared using AutoDock 4.2. Hydrogen atoms were added and the water molecules were removed using the AutoDock Tools (ADT) module included in AutoDock. Charges were adjusted using AutoDock’s Gasteiger charges module for proteins, and atom type was modified to ADT type for calculations. In our calculations, we dock the ligand (natural or *ortho-*vanillin) with ligand-free LasR or RhlR. For each type of atom in the ligand being docked, AutoDock needs a pre-calculated grid map. These maps are calculated using AutoGrid. The Gasteiger-Marsili method was used to determine the atomic charges of the protein. The AutoGrid application created mass-centered grid maps with 80 grid points in each direction and 0.375 spacing. Ten different docking runs for the ligand were carried out, followed by the evaluation of docking results for the binding mechanism and conserved interactions, such as hydrogen bonds and hydrophobic interactions, between the hits and the LasR or RhlR binding site. The common interactions of the ligand-docked complexes were analyzed and the one with the best binding score based on the binding free energy was reported.

### Statistical analyses

All statistical analyses (one-way anova, students t-test) were done using Prism v10. IC_50_ and EC_50_ curves were fitted using a nonlinear regression model with a variable slope unless otherwise stated.

## ACKNOWLEDGEMENTS

This work was supported by the NIH through grant R35GM133572 to J.R.C and R35 GM131817 to H.E.B and by Inez Jay Fund to J.R.C. V.D.C. was supported by an Undergraduate Research Award from the KU Center for Undergraduate Research and a K-INBRE fellowship (P20 GM103418). K.A.T. was supported by KU Center for Undergraduate research Emerging Scholars program, U.S. Department of Education McNair Scholars Program, and the NIH Maximizing Access to Research Careers program (MARC) (T34GM136453-01). V.D.C. was supported by an Undergraduate Research Award from the KU Center for Undergraduate Research and a K-INBRE fellowship (P20 GM103418). R.G.A. was supported by the Fulbright Foreign Student Program (15160174).

