## Supplemental for "Characterization of natural product inhibitors of quorum sensing in *Pseudomonas aeruginosa* reveals competitive inhibition of RhlR by *ortho*-vanillin"

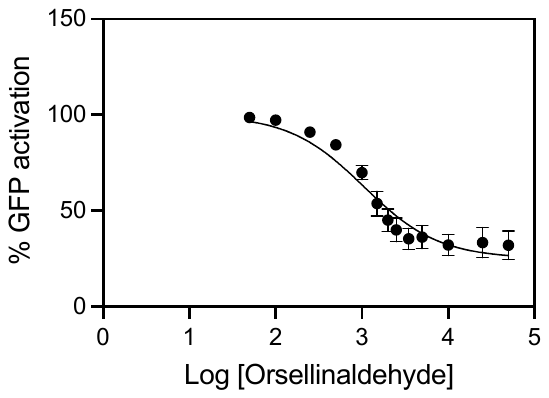

**Fig. S1.** **Orsellinaldehyde inhibition of a constitutive GFP reporter in *E. coli*.** Orsellinaldehyde was used to treat strain *E. coli* carrying pUC18T-mini-Tn7T-Gm-gfpmut3 carrying *gfp* fused to the constitutive *rrnB* promoter similar to the *lacZ* reporter assays described in the Materials and Methods of the main text. The calculated LD_50_ of orsellinaldehyde with GFP was 1057 μM. Results are the averages of three independent experiments and the error bars represent the standard deviation.

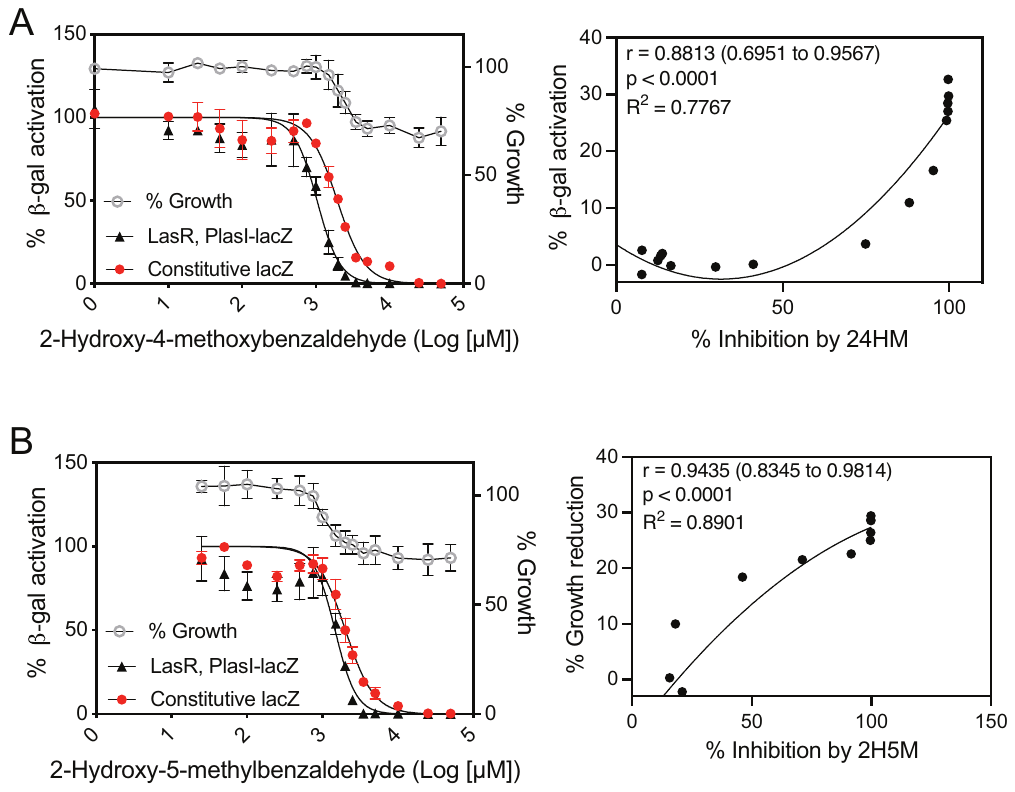

**Fig. S2.** **Activities of 2-hydroxy-4-methoxybenzaldehyde (A) and 2-hydroxy-5-methylbenzaldehyde (B) with LasR and constitutive reporters.** Left column shows growth (grey line) as a percent of *E. coli* cells with no compound and dose-response curves of each compound in competition with 100 nM 3OC12-HSL in cultures of *E. coli* with arabinose-inducible LasR and a LasR-dependent *lasI-lacZ* reporter (black line) or of *E. coli* with a constitutive *aph-lacZ* reporter (red line). IC_50_ values from the fit curves are given in Table 1 in the main text. Results show the averages of three (2H4M) or four (2H5M) independent experiments and the error bars represent the standard deviation. Right column shows average values from the graphs on the left (% growth reduction vs. % inhibition of the *lasI-lacZ* reporter), which were used to determine Pearson’s correlation coefficient (r value) and significance (p) and generate a fitted line using a simple linear regression model (B) or a second-order polynomial nonlinear regression model (A).

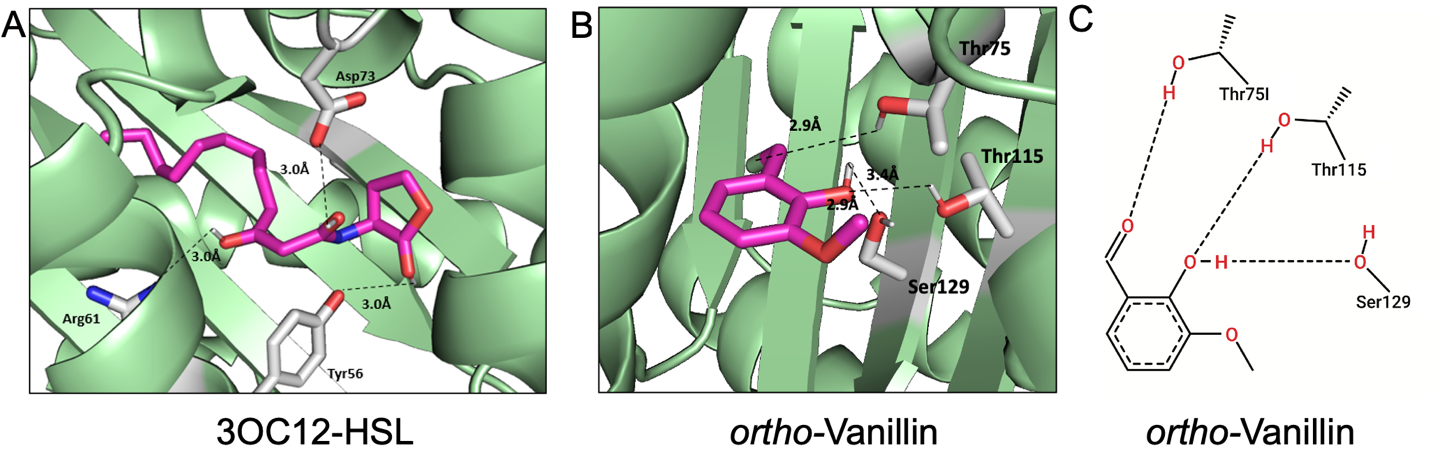

**Fig. S3. Molecular docking of *ortho*-vanillin in the LasR binding site** (PDB ID:6V7X): (A and B) 3D representation of the native ligand 3OC12-HSL or *ortho*-vanillin (magenta sticks) in the LasR binding site (green ribbons with key residues shown in sticks). Dashed lines indicate predicted polar interactions between the ligand and the LasR amino acid residues (Asp73, Tyr56 and Arg61 for 3OC12-HSL and Thr75, Thr115 and Ser129 for *ortho-*vanillin). The compound was also predicted to have interactions within 4 Å with the LasR residues Leu110, Trp88, Ala105, Tyr64, Leu36, Tyr56, Trp60, and Gly126. (C) 2D interaction map of *ortho-*vanillin and Thr75, Thr115 and Ser129 with dashed lines representing hydrogen bond interactions. The docking score was -4.51 kcal/mol for *ortho*-vanillin, and -6.79 kcal/mol for 3OC12-HSL.

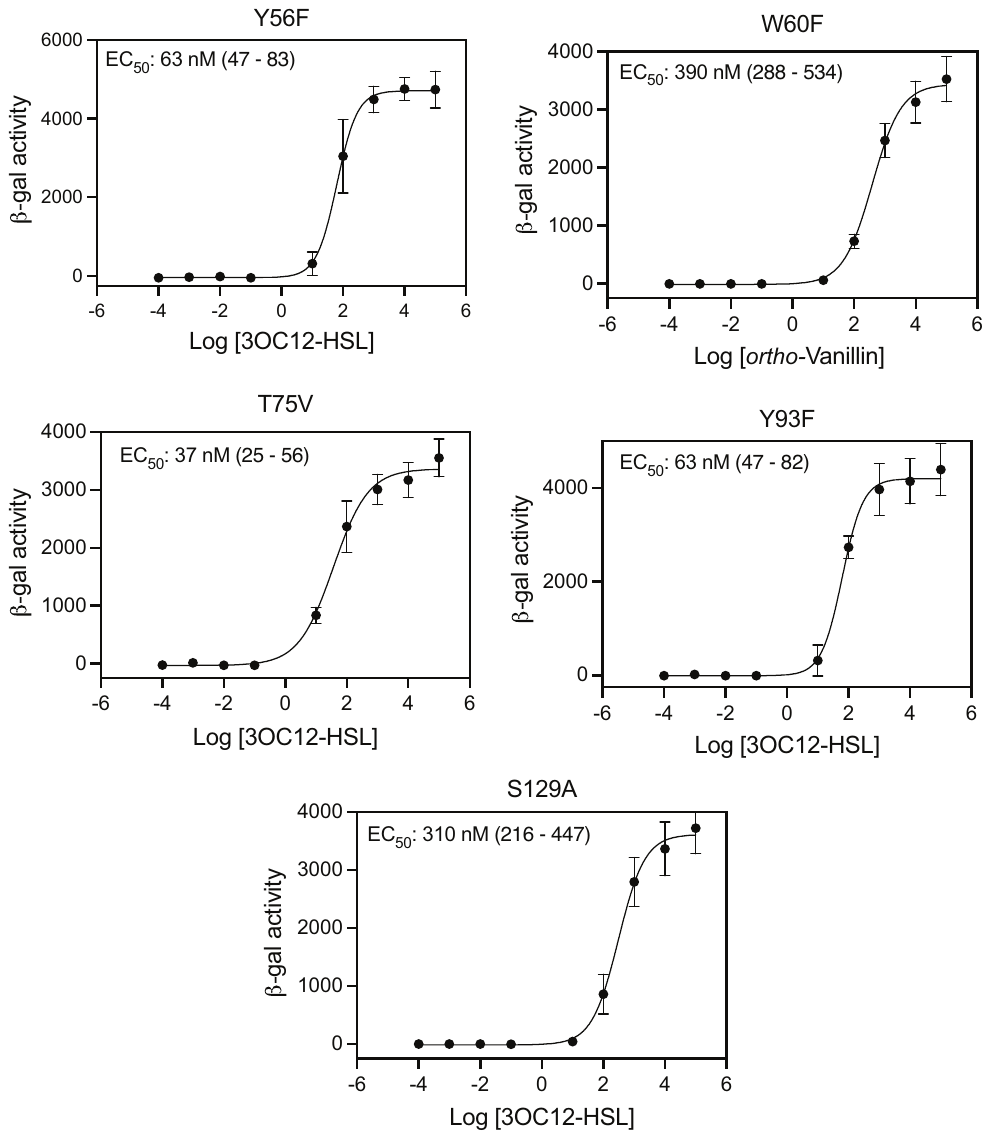

**Figure S4. Dose-response agonism data for 3OC12-HSL and LasR mutants.** Experiments were conducted with *E. coli* carrying pJN105-L (encoding arabinose-inducible *lasR*) and pSC111-L (encoding the LasR-dependent *lasI-lacZ* reporter) or a pSC111-L derivative encoding the indicated LasR mutant. In each case the curve was fitted using a nonlinear regression 4-parameter (variable slope) model in Prism v10. IC_50_ values from the fit curves are given within each panel with the 95% confidence interval indicated in parentheses. Data show the averages of four independent experiments and the error bars represent the standard deviation. Results were used to determine the concentration of 3OC12-HSL to use for LasR mutant dose-response curves with *ortho-*vanillin (Fig. 5 in the main text).

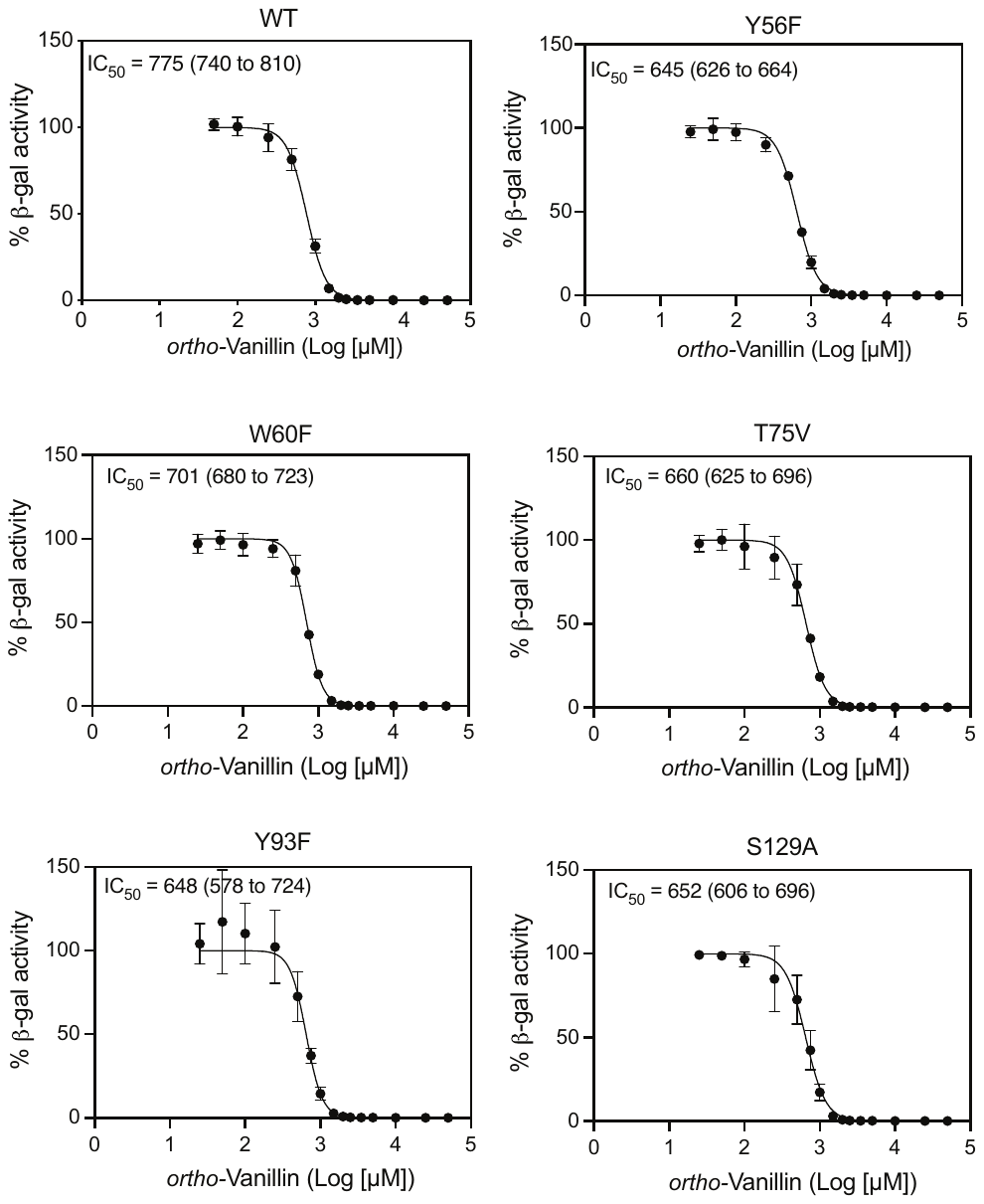

**Fig. S5. *ortho-*Vanillin antagonism data for LasR mutants.** Experiments were conducted with *E. coli* carrying pJN105-L (encoding arabinose-inducible *lasR*) and pSC111-L (encoding the LasR-dependent *lasI-lacZ* reporter) or a pSC111-L LasR-mutant derivative. Dose-response experiments with *ortho-*vanillin were carried out by competing with the EC_50_ of 3OC12-HSL determined for each LasR mutant from Fig. S5. IC_50_ values from the fit curves are given within each panel with the 95% confidence interval indicated in parentheses. Results show the averages of four independent experiments and the error bars represent the standard deviations. The generated IC_50_ data from these curves were used to plot the values in Fig. 5A in the main text.

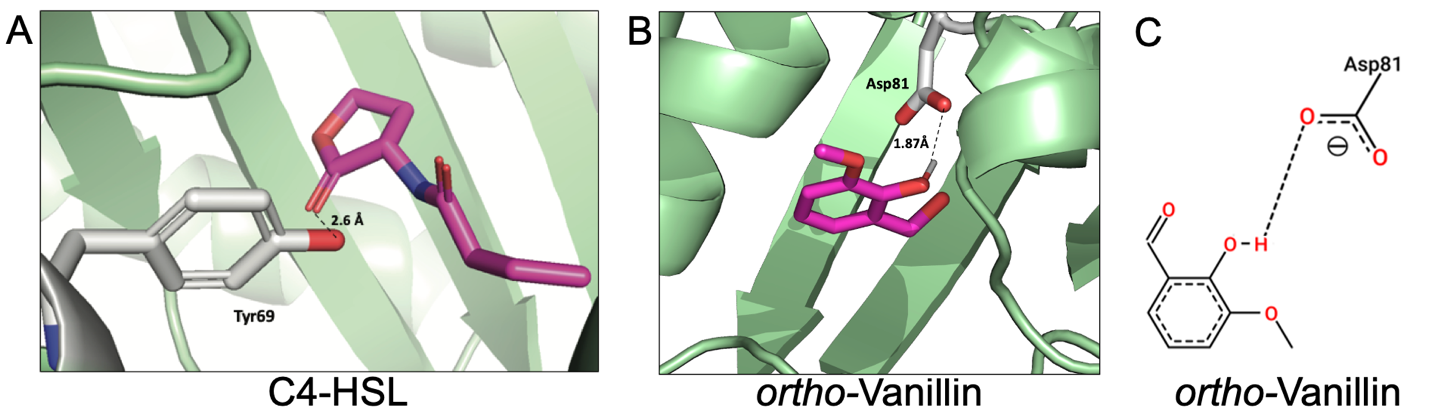

**Fig. S6. Molecular docking of *ortho*-vanillin in the RhlR binding site** (PDB ID:8DQ0): (A and B) 3D representation of the native ligand C4-HSL *ortho*-vanillin (magenta sticks) in the RhlR binding site (green ribbons with key residues shown in sticks). Dashed lines indicate predicted polar interactions between each compound and RhlR (Tyr69 for C4-HSL and Asp81 for *ortho-*vanillin). C4-HSL was also predicted to have interactions within 4 Å with the RhlR residues Trp93 and Ala108. (C) 2D interaction map showing the hydrogen bond between *ortho*-vanillin and the RhlR residue Asp81. The docking score was -5.20 kcal/mol for *ortho-*vanillin, and -5.21 kcal/mol for C4-HSL.

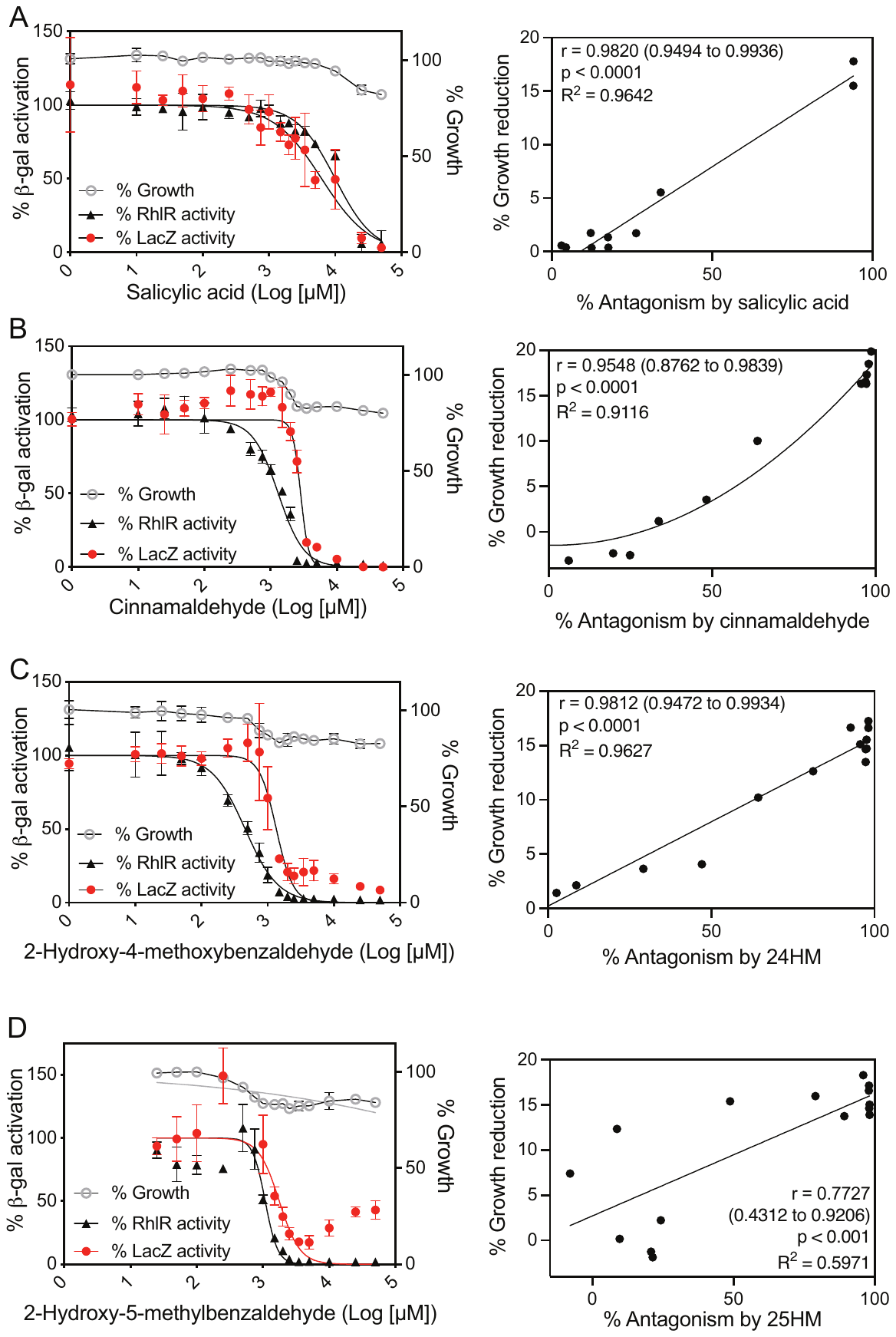

**Fig. S7. Activities of compounds with RhlR and constitutive reporters.** Left column shows growth (grey line) as a percent of E. coli cells with no compound and dose-response curves of each compound in competition with 100 nM 3OC12-HSL in cultures of *E. coli* with arabinose-inducible LasR and a LasR-dependent *lasI-lacZ* reporter (black line) or of *E. coli* with a constitutive *aph-lacZ* reporter (red line). In each case the curve was fitted using a nonlinear regression 3-parameter model. Curves were fitted using a nonlinear regression 3-parameter model with the top and bottom constrained to 100% and 0% respectively for bioreporters or with the top only constrained for growth. IC_50_ values from the fit curves are given in Table 1. Results show the averages of three independent experiments and the error bars represent the standard deviations. The right column shows average values from the graphs on the left (% growth reduction vs. % inhibition of the lasI-lacZ reporter), which were used to determine Pearson’s correlation coefficient (r value) and significance (p) and generate a fitted line using a simple linear regression model (B) or a second-order polynomial nonlinear regression model (A and C).

**Table S1. Bacterial strains and plasmids used in this study.**

| Strain or plasmid | Relevant properties^a^ | Reference or source |
| --- | --- | --- |
| *Strain* |  |  |
| *E. coli* DH5α | [F- φ80 lacZΔM15] Δ(lacZYA-argF)U169 recA1  endA1 hsdR17, [rK−mK+] supE44 thi-1 gyrA relA1 | Invitrogen |
| *Plasmids* |  |  |
| pSC11-L | Broad host range *lasI-lacZ* reporter; Ap^R^ | (1) |
| pJN105-L | Arabinose-inducible *lasR* expression vector; Gm^R^ | (2) |
| pECP61.5 | Vector for IPTG-inducible *rhlR* expression and a *rhlA-lacZ* reporter; Ap^R^ | (3) |
| pVT19 | Vector with constitutive *aph-lacZ* reporter | This study |
| pUC18T-mini-Tn7T-Gm-gfpmut3 | Vector with constitutive *rrnB-gfp* | (4) |
| pJG044 | Y56F mutant analog of pJN105L | (5) |
| pJG008 | W60F mutant analog of pJN105L | (5) |
| pJG011 | T75V mutant analog of pJN105L | (5) |
| pJG012 | Y93F mutant analog of pJN105L | (5) |
| pJG014 | S129A mutant analog of pJN105L | (5) |

^a^Abbreviations: Ap^R^ = Ampicillin resistance; Gm^R^ = Gentamicin resistance.
